# European freshwater macroinvertebrate richness and abundance: alternative analyses and new findings

**DOI:** 10.1101/2024.08.13.607735

**Authors:** Benoît O.L. Demars

**Affiliations:** Norwegian Institute for Water Research (NIVA), Økernveien 94, 0579 Oslo, Norway

**Keywords:** macroinvertebrates, time series, river, richness anomaly, taxonomic resolution, sampling effort

## Abstract

1. Studies at local to national extent have documented a recovery in macroinvertebrate taxonomic richness following improvements in water quality. The study by Haase *et al*. (2023) published in Nature claimed that the overall recovery came to a halt across Europe by 2010. However, the lack of monitoring design, the varying lengths in time series and heterogeneous taxonomic resolution (species, genus and families), along with insufficient information on data handling prior to statistical analyses are raising questions about the reliability of the findings.
2. Here I use the open access raw data of the original study to calculate the proportion of taxa identified to the targeted taxonomic resolution (species, genus or family), which revealed a lack of taxonomic consistency within some of the time-series. I then devised a simple taxonomic correction to remove potential biases in the richness trend estimates through the modelling procedures using linear models.
3. In order to make the data more comparable across studies and over time, I calculated an anomaly in taxonomic richness relative to a five-year reference period within 1990-2020, so all time series (≥15 years long, ≥8 samples) overlapped. The concept is borrowed from the familiar temperature anomaly in climate research to track deviations from a norm. I ran non-linear trend analyses to reveal changes in the anomaly in taxonomic richness during the period 1990-2020.
4. European taxonomic richness using 1816 sites in 47 studies (full dataset) increased linearly by about 0.29±0.09 taxa per year when using all taxonomic ranks (species, genus, family), compared to the average 0.20 taxa per year in the original study, but dropped to 0.15±0.04 taxa per year at family level. The same results were produced after geographical thinning to 687 sites separated by at least 20 km from each other’s. Further data analyses revealed the extent of discrepancies in taxonomic resolution (proportion of taxa identified to species or genus level) within time-series and its impact on trend estimates.
5. The linear increase in abundance over time was marginal (1 individual / year or 0.12% of average abundance) in the full dataset and not significant within 1990-2020 period, contrary to published findings (1.17%) due to a calculation error in the original study.
6. The linear analyses of species richness were run on centred years and did not allow the study of the temporal dynamics in taxonomic richness. Non-linear analyses using the anomaly in taxonomic richness for the period 1990-2020 revealed no change in taxonomic richness apart from a post millennium small and short rise using all taxonomic ranks (1120 sites, 27 studies), possibly due to a concurrent increase in sampling effort (abundance) across sites.
7. Coarsening the taxonomic resolution to family level did not alter the dynamic of the anomaly in taxonomic richness over time, possibly a result from poor sample sampling effort. The average ‘species’ richness (762 sites) was about 30 taxa per sample, barely higher than family richness (20 taxa per sample) and very small compared to studies with more intensive sampling efforts. Independently of the effect of anthropogenic impacts, I question the adequacy of the current biomonitoring design and sample sampling effort to study river macroinvertebrate biodiversity.
8. *Implications of new findings*. Linear trend estimates in taxonomic richness (independently of the time period) were dependent on taxonomic resolution, higher at ‘species’ than family level. Neither the abundance nor the anomaly in taxonomic richness showed signs of recovery during the period 1990-2020. Current sampling efforts for rapid bioindicators, such as those developed for the European Water Framework Directive, are inadequate to address the needs of the EU 2030 Biodiversity Strategy. Macroinvertebrates would be right to demand more from us.

## 1. Introduction

European rivers are the recipients of rural, urban and industrial effluents, as well as diffuse pollution from land use and atmospheric deposition. Rivers have also been physically transformed: straightened, deepened and dammed losing their natural connectivity and altering the flow of water and sediment. The impact of gross pollution on river macroinvertebrates and fish has been well documented (Carpenter, 1928; Hynes, 1960). The stench of gross pollution, the lack of fish and environmental movements pushed successive governments across Europe into adopting new legislation to curb pollution with the aim to improve sanitation and fisheries: early national legislations included the UK Rivers Pollution Prevention Act of 1876, and UK Salmon and Freshwater Fisheries Act of 1923), before increasingly stringent EU regulations were adopted (e.g. Urban Waste Water Treatment Directive 91/271/EEC, EU Water Framework Directive 2000/60/EC). The ecological quality of rivers has improved following the removal of gross pollution (Langford *et al*., 2009; Whelan *et al*., 2022).

National studies have also documented recoveries of macroinvertebrate biodiversity with improvements in water quality (Vaughan & Ormerod, 2012; Vaughan & Ormerod, 2014; Qu *et al*., 2023). In England and Wales, the recovery of family richness reached a plateau or slowed considerably at the turn of the millenium (Vaughan & Ormerod, 2012; Qu *et al*., 2023).

It remains difficult, however, to estimate the effectiveness of new directives and policies across Europe(e.g. UWWTD, WFD, EU Biodiversity Strategy for 2030). Peter Haase and Ellen Welti have directed a team of 95 co-authors to provide long time series of river macroinvertebrates across Europe, with the aim of testing for a Europe-wide recovery in the biodiversity of river macroinvertebrates (Haase *et al*., 2023). While the aim is simple, the lack of overall sampling design, time series of various lengths not necessarily continuous or overlapping in time, variability in sampling methods and taxonomic resolution renders the task challenging. Haase *et al*. (2023) established six key criteria to select the data: “1) inclusion of abundance estimates, 2) surveyed whole freshwater invertebrate communities (not restricted to certain taxonomic groups, such as insects), 3) identified most major taxa to family, genus or species, 4) had a minimum of eight sampling years (not necessarily consecutive), 5) had no changes in sampling method or taxonomic resolution during the sampling period, and 6) had consistent sampling effort per site (e.g. number of samples or area of river sampled) across years”.

Haase *et al*. (2023) reported an average linear increase in taxonomic richness of 0.20 taxon per year across all 1816 time series in 47 studies from 22 European countries. The results from a Bayesian meta-analysis of trend estimates derived from linear models (two step procedure) or linear mixed effect models (one step procedure) both gave similar results. These linear models included an autoregressive term to deal with temporal autocorrelation. Study and country were used as random factors which likely lessened potential (untested) issues with spatial autocorrelation. The year of sampling was also centred to improve convergence of the model, and together with the use of linear models, prevented the detection of possible variability in the overall trend estimate over several decades. Haase *et al*. (2023) devised a 10-year moving-window analysis over the period 1990-2020 (using the same modelling approach as above) to claim that trend estimates in taxonomic richness were increasing (by about 0.27 taxa per year) in the 1990s and 2000s but reached a plateau from 2010 onwards. The authors also quantified other aspects of biodiversity and tried to assign possible causes for the observed differences in trend estimates across Europe, but here I will focus only on the trend estimates in taxonomic richness and abundance, which were the most striking results in Haase *et al*. (2023) given the title of their paper: “The recovery of European freshwater biodiversity has come to a halt”.

This is, thus far, the most comprehensive study of macroinvertebrate time series across Europe and is a major achievement. Yet, the results are a product of the data at hand (i.e. they reflect responses to local conditions and non-random site selection, e.g. Jähnig *et al*., 2021), and are not representative of Europe, or its habitat loss and associated biodiversity crisis (e.g. Albert *et al*., 2021). This is a limitation common to work on time series at large spatial extent (e.g. van Klink *et al*., 2020; Blowes *et al*., 2019), due to the lack of a continental or global monitoring design for biodiversity.

Haase *et al*., 2023 were fully aware of key limitations in their approach and strove to convince the reader through the use of sensitivity analyses that their results were robust. Nonetheless, at least two major caveats remain. The first one is that the time series do not or only partially overlap in time, so the change in trend estimate may simply be due to an artefact of site selection; in the words of Haase *et al*. (2023): “These data were not selected randomly but were collected from available studies that met our six criteria. As these data were collected from sites exposed to varying and unquantified levels of anthropogenic impacts, we cannot rule out biases arising from unequal representation of sites exposed to different impact levels from severely impacted to least impacted.”; “We cannot fully discount the possibility that biases in the characteristics of sites sampled across time affected trajectory results.” The second caveat is that those time series may also have different taxonomic resolution. The authors grouped their data into ‘species’ (the quotes are mine), genus (mixed) and family levels, but the ‘species’ data are actually mixed-taxon data and the taxonomic resolution is likely to differ substantially between the individual researchers or consultants that carried out the identification (Haase *et al*., 2010; Haase *et al*., 2006). Previous studies on river macroinvertebrate biodiversity either used family level (consistency in taxonomic resolution, e.g. Vaughan & Ormerod, 2012; Vaughan & Ormerod, 2014) or species level data collected by taxonomists (e.g. Domisch *et al*., 2013).

The selection of such heterogeneous data for time series analyses is challenging, and yet I could not find any information on how the data were cleaned. What taxa were removed from the database used (e.g. higher taxonomic rank than families)? Were only species used or were genus, family and higher taxa groups within the ‘species’ data kept? Were larvae and adults, often recorded separately (common in coleoptera), aggregated? How was double accounting prevented, e.g. when a taxon is a genus with a single species but not always identified down to species, were they aggregated? The data were recently published in a more accessible form, but the authors did not provide additional information (Welti *et al*., 2024).

Finally, I found it personally difficult to relate to the results presented: the 0.73% and 1.17% increase in taxon richness and abundance, respectively, did not tell me how many taxa the rivers of Europe have gained over a year or a decade. The reader had to fetch a table provided in GitHub (equationsToPercChangePerYr.xlsx) to recover the trend estimate as 0.20 taxa year^-1^, equivalent to two taxa per decade for an estimated average richness of 27 taxa on the centred year. The percentage increase in trend estimate for abundance did not appear to be correct (see Discussion).

Thanks to the authors for providing the raw data used in their analyses, I had the opportunity to explore alternative approaches. I therefore propose in this study to highlight some properties of the data, including taxonomic issues, not presented by Haase *et al*. (2023). Then, I will propose a solution to deal with, rather than ignore, taxonomic heterogeneity and explore the potential impact on the overall trend estimate results on centred year (0.20 taxa per year). I will also propose an alternative method of the moving window analysis by using the anomaly in taxonomic richness against a reference period. The concept is borrowed from the familiar temperature anomaly in climate research to track deviations from a norm. I will apply corrections for the heterogeneity in taxonomic resolution in individual time series. I will also repeat my analyses after converting taxonomic resolution to family level. My analyses were designed to be spatially explicit to look for general trends in richness and abundance (*cf* Pilotto *et al*., 2020) and lessen the biases arising from unequal representation of sites through time, using long time series with at least a partial overlap, while keeping as many sites as possible.

## 2. Methods

### 2.1. Data management

First I recompiled a single file from the raw taxonomic composition data for all included studies, which held 727696 records with the site id, year, individual taxa names with their abundance and taxon id from Freshwaterecology.info (https://github.com/Ewelti/EuroAquaticMacroInverts, Raw Data/csvs_each_dataset folder). I had to make minor changes to few individual files to make sure all files had the same number of columns to be automatically compiled in R version 4.3.2 and write the file df_compiled.txt. I then extracted, in Excel (Microsoft), a list of 2666 taxa and scored each taxon with a taxonomic rank: species (s), genus (g), family (f) and higher taxa (x). I extracted a full list of taxa with taxon id, TaxaGroup, family and species names held by www.freshwaterecology.info - the taxa and autecology database for freshwater organisms, version 8.0 (accessed on 24.05.2024) – Schmidt-Kloiber and Hering (2015). The genus names were extracted from the species names. I added manually the name of taxonomic rank for 287 taxa listed in Haase *et al*. (2023) but not yet included in the open access freshwaterecology.info database. I added a taxon id missing for Platyhelminthes, Nemertea, Mollusca and Clitellata. I then appended the taxonomic rank and their names to the full list of individual records and saved the file (df_compiled_taxa_rank.txt). I used R for the remaining data handling. I deleted three rows with abundance = 0 and removed taxa identified to taxonomic rank higher than family level (x). I found several issues with the sample_id column, so I concatenated site_id and year instead (code). The data held 712786 records, 26664 samples, in 1816 sites and 47 studies.

I then calculated species, genus and family richness for all samples, and the average per site. Species richness corresponded to the highest taxonomic resolution and included taxa identified to species, genus and family – correspond to spp_richness calculated by Haase *et al*. (2023). The genus richness included genus and family (species names were replaced by their genus name). The family richness included all species and genus names replaced by family names. This approach is different from Haase *et al*. (2023): all species names are here nested within genus and family names; this is more accurate than the original study which assigned ‘species’, ‘genus’ or ‘family’ level of identification for whole samples and time series, when in fact each sample had macroinvertebrate individuals identified to mixed taxa levels. Further, the distinction of taxonomic resolution (s, g, f) at the level of the individual records allowed the calculation of the proportion of taxa identified to species (Fsp), genus (Fg), mixed taxa (species and genus, Fm) and family (Ff) for each sample (range 0 to 1). I also estimated the fraction of taxa identified to the highest resolution (Fy) assigned by Haase *et al*. (2023) using Fsp for ‘species’, Fm for ‘genus’ and Ff for ‘family’. I centred all these proportions around their median within each time series (c.Fsp, c.Fg, c.Fm, c.Ff, c.Fy), range -1 to 1 (see Fig. 1). I rounded those values to one decimal so I could use them as factor in statistical analyses.

**Fig. 1.**
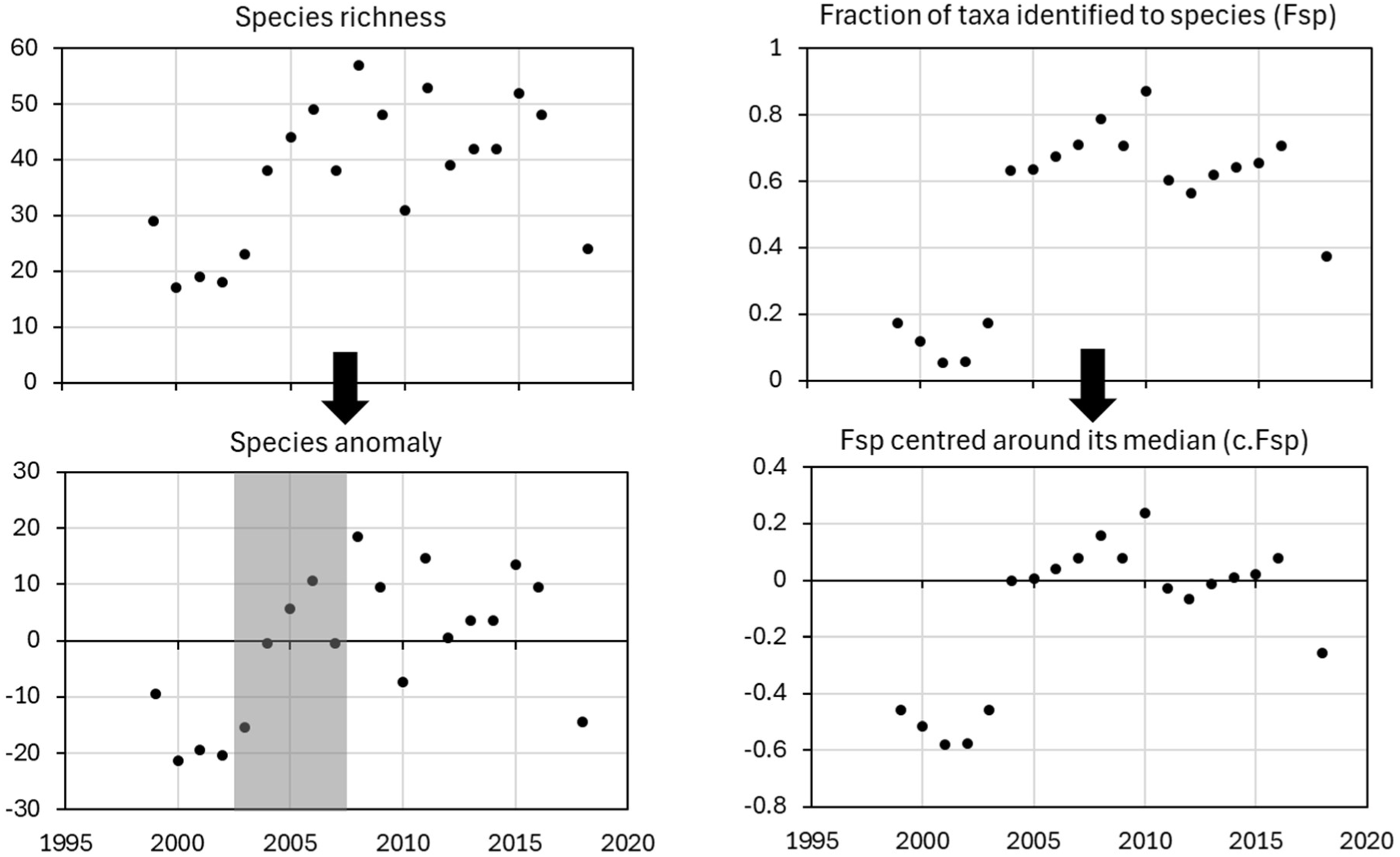
Data transformation for the first site of the Danish study (site 107000001). Left panels: Species richness anomaly relative to average species richness in the reference period (2003-2007) highlighted in grey. Right panels: proportion of taxa identified to species level centred around its median.

I then opened (in R) the file containing data for each site and year, and taxonomic diversity metrics All_indices_benthicMacroInverts_AllYears_alienzeros.csv (https://github.com/Ewelti/EuroAquaticMacroInverts, Outputs folder). I recalculated the centred year (cYear) and year order (iYear) as in Haase *et al*. (2023), before I deleted rows without records. I appended latitude and longitude from AquaticMacroInvert_metadata.xlsx (https://github.com/Ewelti/EuroAquaticMacroInverts). I then added the species, genus and family richness, as well as all the proportion of taxa identified to specific taxonomic ranks calculated above. The file was saved as Haase_summary_taxa_rank.txt.

Finally, I produced two data subsets for additional statistical analyses. I calculated the average taxonomic richness over the period 2003-2007 for each time series, from which I derived a species richness anomaly for each times series (sample richness – average richness) – see Fig. 1). Thus, taxonomic richness becomes comparable across time series (period 1968-2020) and at least all time series overlapped (91% of the samples, 88% of the sites were included). Since there were too few sites in the early years, I then retained time series with a minimum range of 15 years and at least 8 samples within the period 1990-2020 (70% of the samples, 62 % of the sites were included). Note, restricting the period to 1992-2019 as in Sinclair *et al*. (2024) produced the same results. This subset had varying taxonomic assignments (as defined by Haase *et al*., 2023) to ‘species’, ‘genus’ (mixed taxa) and ‘family’, so I also produced another subset following the same rules but using only the ‘species’ times series (38% records, 23% sites were included). The R code is available in GitHub (Taxonomic_richness_anomaly.R).

Since richness is generally highly dependent on sampling effort, I calculated a site level abundance anomaly by subtracting the intra-site median abundance to the sample abundance (including all taxa, as reported in the original study), and rescaled to positive values to get the same lowest number of individuals per sample as for abundance. This rescaling allowed a necessary logarithmic transformation for the statistical analyses. Few abundance outliers (on the logarithmic scale) were removed prior to data analyses: number of individuals exceeding 1,000,000 (one sample) and 100,000 (three samples) for all taxonomic ranks and ‘species’ data, respectively, for the period 1990-2020.

### 2.2. Statistical analyses

#### Trend estimates

I used linear mixed effects models (nlme R package version 3.1-164, Pinheiro *et al*., 2023; Pinheiro & Bates, 2000) to estimate trends in taxonomic richness against centred year (cYear) as fixed effect and site as random effects. The temporal autocorrelation was at most +0.11 at one year lag and -0.17 at five year lag, and the autoregressive term (AR1) did not improve much the (still significant) autocorrelation in model residuals (Fig. S1). The same results were obtained when taxonomic richness was corrected prior to running the model. Thus, I did not use a temporal autoregressive term in subsequent analyses (R script available in GitHub, Haase_nlme.R). I tested the response of taxonomic richness or abundance against cYear as fixed effect and site or/and study with/without a correction for taxonomic resolution (c.Fy for all taxa or c.Fsp for species) as random factors, with the lme4 R package (version 1.1-35.1, Bates *et al*., 2015b; Bates *et al*., 2015a; R script examples in GitHub, Haase_lme.R). I did not include country as a random effect since it had completely negligible effects once study was selected. I used linear models to test if the correction for taxonomic resolution (c.Fy) altered the parameters of the model (slope and intercept). I also plotted the distribution of c.Fy to highlight the number of samples suffering from taxonomic discrepancy within time series. Note c.Fy was rounded to one decimal point and used as a factor in all trend estimate analyses. Finally, spatial autocorrelation (spline correlogram) in the site trend estimates (*not the model residuals*) of selected linear mixed effect models was estimated with the ncf R package (version 1.3-2, Bjornstad, 2022), after transforming the WGS84 coordinates into the projected coordinate system for Europe (EPSG:3035) using the sf R package (version 1.0-15, Pebesma, 2018; Pebesma & Bivand, 2023; R script examples in GitHub, lme_autocorrelation_test.R). I then re-run linear mixed effects models after thinning the dataset geographically, based on the results of the trend estimate autocorrelation, using spThin R package (Claessens, Kyritsis & Atkinson, 2023; R script in GitHub, Haase_lme_thinned.R). Abundance was transformed with a natural log prior to running the models to improve homoscedasticity and normality. All models assumed normally distributed errors, which were checked visually using histograms for both fixed and random effects (intercept and slope).

#### Taxonomic richness anomalies

I used spatial gam to test the effects of year and site location (latitude/longitude), with/without taxonomic correction (c.Fy for all taxa or c.Fsp for species) on the response variables abundance or taxonomic richness using mgcv R package (version 1.9-1, Wood, 2017; R script examples in GitHub, Haase_gam.R). Note the taxonomic corrections (c.Fy, c.Fsp) were used as numeric values, except for ‘family’ where the correction (c.Fm) was not needed. The effects of study and country were negligible and not included in the spatial models. Abundance and abundance anomaly were transformed with a natural log prior to running the models to improve homoscedasticity and normality. General additive model uses non-parametric functions (so-called splines) able to model non-linear changes in taxonomic richness against time. I used reduced ranked thin plate regression spline basis, with automatic selection of the effective degrees of freedom for all smooths (Wood, 2003; Wood, 2017). The smoothness selection was by REML. I plotted partial effect plots to show the prediction (from the fitted GAMs) of the selected co-variable (year), setting other predictive variables at their average values. Standard errors on the plots show the 95% confidence interval. I checked the model converged rapidly towards a solution, the number of nodes for the splines was appropriate, concurvity of individual and pairwise variables (equivalent to collinearity in linear models), and the residuals of the model were normally distributed. Spatial autocorrelation (spline correlogram) in the response residuals (anomaly in taxonomic richness) of selected years in the gam models was estimated with the ncf R package, as above. Alternative gamm models with taxonomic correction and study as random factors gave virtually the same results (not shown but R script examples in GitHub, Haase_gamm.R). Additionally, gamm models following the same approach as Sinclair *et al*. (2024) showed some significant temporal autocorrelation but the results were virtually unchanged with or without an autoregressive term (not shown but R script examples in GitHub, Haase_gamm.R). Finally, the fitted anomaly in taxonomic richness was regressed against fitted abundance for the period 1990-2020 to test the potential effect of sampling effort across sites. The dataset was thinned to odd years to remove or lessen temporal autocorrelation of model residuals. Temporal autocorrelation was tested with the Breusch-Godfrey test (third-order serial correlation) included in the lmtest R package (Zeileis & Hothorn, 2002; but R script examples in GitHub, Haase_gam fit regression with autocorrelation).

## 3. Results

### 3.1. Quick look at the data

The first striking issue is that median (and interquartile) taxonomic richness from 1968 to 1979 remained below 10 taxa with maximum values below 25 taxa per sample (Fig. 2). Taxonomic richness was around 10 taxa per sample in the 1980s with maximum values generally below 50 taxa. The number of sites per year was low (<100) until the early 1990s and peaked during 2003-2016 with well over 1000 sites per year (Fig. 3). The proportion of taxa identified to species, genus and families changed considerably over the five decades. Identification to species level was notably low in the 1990s.

**Fig. 2.**
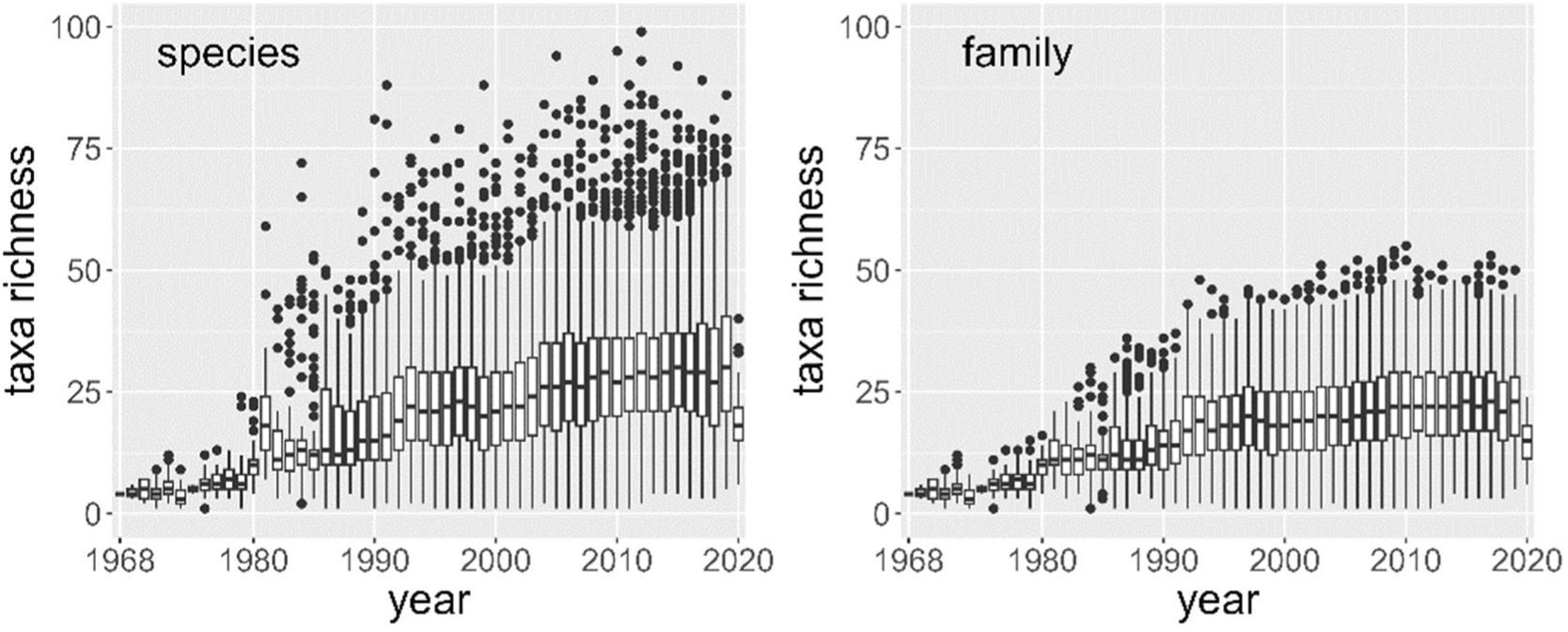
Annual changes in taxonomic richness over five decades at 1816 sites with two levels of taxonomic resolution (‘species’ and family). Boxplots represent the median and interquartile, whiskers follow the rule 1.5 x interquartile range, and dots represent outliers.

**Fig. 3.**
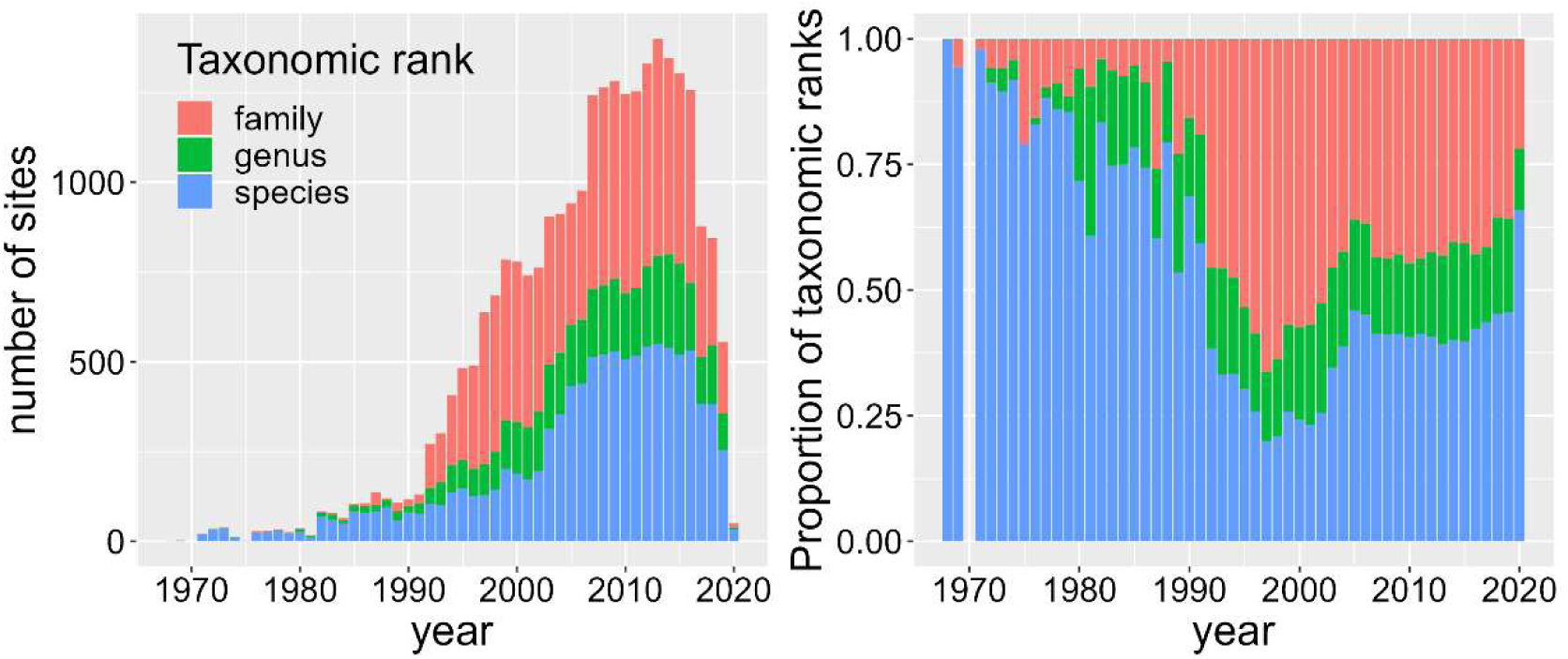
Annual changes in the number of sites over five decades and sum of proportion of taxa per sample (n=26664) identified to species (Fsp), genus (Fg) and family (Ff) level at 1816 sites.

### 3.2. Trend estimate analyses using centred year

The trend estimates of taxonomic richness per year using all 1816 sites in 47 studies ranged between 0.26±0.09 and 0.29±0.09 with or without correction (Table 1). Similar results were produced on the subset of ‘species’ data including 762 sites in 28 studies. However, the taxonomic correction lowered significantly the intra-site trend estimate by 31% from 0.32±0.02 to 0.22±0.03 taxa year^-1^ (without study as random factor). Intra-site inconsistencies in taxonomic resolution had a large effect on both the trend estimate (slope) and the average taxonomic richness (intercept) – Fig. 4. The proportion of intra-site taxonomic discrepancies (centred Fy) exceeding ±0.3 revealed 907 samples with poor and 72 samples with ‘overzealous’ taxonomic determination (Fig. 4). Trend estimates were much lower 0.15±0.03 and 0.17±0.03 when all taxonomic ranks were converted to family level for 1816 sites in 47 studies and 762 sites in 28 studies, respectively (Table 1).

**Fig. 4.**
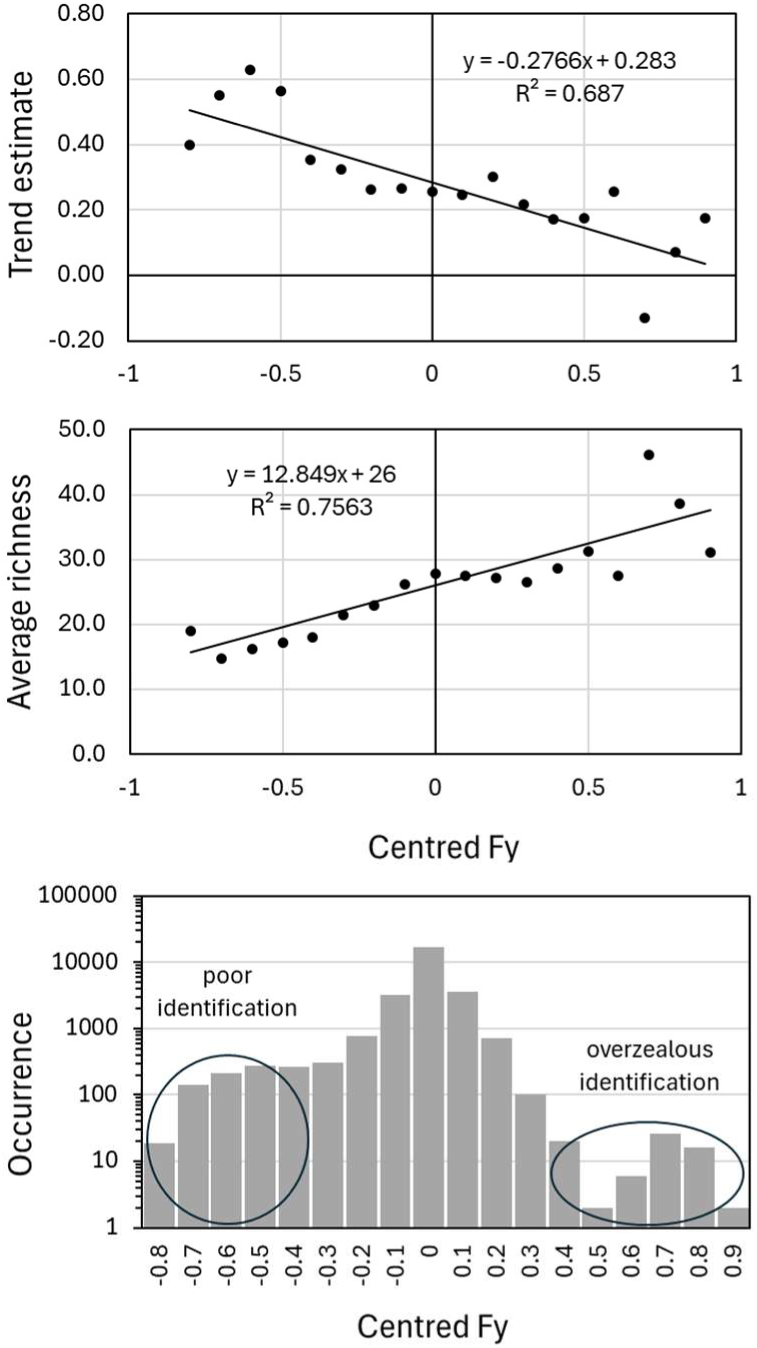
Effect of intra-site discrepancy in taxonomic resolution on trend estimate (slope) and average richness (intercept). Taxonomic resolution was quantified as the proportion of taxa (Fy, range 0-1) identified to species (Fsp), mixed taxa (species + genus, Fm) or family (Ff) within a sample. Fy was then centred (c.Fy) within individual time series. Centred Fy tends to zero where taxonomic resolution was consistent within sites (individual time series). Centred Fy below or above 0.3 denote samples with poor identification (907 cases) or ‘overzealous’ identification (72 cases), respectively, out of 26664 samples.

**Table 1.** Effects of linear mixed-effects models on average taxonomic richness (intercept) and trend estimate (slope). The response term Species_richness refers to the highest taxonomic resolution available in the dataset and include species, genus and families (even in the ‘species’ data). All taxonomic names were converted to family level and the analyses repeated.

| Lme models | Intercept (se) | Slope (se) | P |
| --- | --- | --- | --- |
| **Including all 1816 sites in 47 studies with different taxonomic resolutions** |  |  |  |
| Species_richness ~ cYear + (1+cYear\|site_id) | 27.1 (0.3) | 0.28 (0.01) | <2.2e<sup>-16</sup> |
| Species_richness ~ cYear + (1+cYear\|site_id) + (1+cYear\|study_id) | 28.6 (1.6) | 0.26 (0.09) | 0.005 |
| Species_richness ~ cYear + (1+cYear\|site_id) + (1+cYear\|c.Fy_id_round) | 26.0 (2.0) | 0.28 (0.05) | 0.0005 |
| Species_richness ~ cYear + (1+cYear\|site_id) + (1+cYear\|study_id) + (1+cYear\|c.Fy_id_round) | 26.3 (2.3) | 0.29 (0.09) | 0.003 |
| **Including all 1816 sites in 47 studies at family taxonomic resolution** |  |  |  |
| Family_richness ~ cYear + (1+cYear\|site_id) | 21.3 (0.2) | 0.20 (0.01) | <2.2e<sup>-16</sup> |
| Family_richness ~ cYear + (1+cYear\|site_id) + (1+cYear\|study_id) | 20.8 (0.9) | 0.15 (0.04) | 0.001 |
| **Including only 762 sites in 28 studies with ‘species’ data** |  |  |  |
| Species_richness ~ cYear + (1+cYear\|site_id) | 27.0 (0.5) | **0.32 (0.02)** | <2.2e<sup>-16</sup> |
| Species_richness ~ cYear + (1+cYear\|site_id) + (1+cYear\|study_id) | 32.0 (2.1) | 0.40 (0.12) | 0.002 |
| Species_richness ~ cYear + (1+cYear\|site_id) + (1+cYear\|c.Fsp_id_round) | 25.8 (2.7) | **0.22 (0.03)** | 1.4e<sup>-05</sup> |
| Species_richness ~ cYear + (1+cYear\|site_id) + (1+cYear\|study_id) + (1+cYear\|c.Fsp_id_round) | 29.5 (3.2) | 0.33 (0.08) | 0.0003 |
| **Including all 762 sites in 28 studies at family taxonomic resolution** |  |  |  |
| Family_richness ~ cYear + (1+cYear\|site_id) | 18.5 (0.3) | **0.16 (0.01)** | <2.2e<sup>-16</sup> |
| Family_richness ~ cYear + (1+cYear\|site_id) + (1+cYear\|study_id) | 21.3 (1.3) | **0.17 (0.03)** | 0.0001 |

The trend estimates were less affected by the taxonomic correction when the data were summarised by studies. Yet, the investigation of random effect coefficients pointed out several studies where the taxonomic correction had a large effect on trend estimates: -0.18 taxa year^-1^ for Denmark (248 sites), +0.42 taxa year^-1^ for Spain_3.1 (121 sites) and -1.01 taxa year^-1^ for UK_4 (12 sites). A quick look at the raw data revealed that these three studies had intra-site inconsistencies in the proportion of taxa identified to species (Denmark, UK) or genus (Spain). This is illustrated for Denmark in Figure 5.

**Fig. 5.**
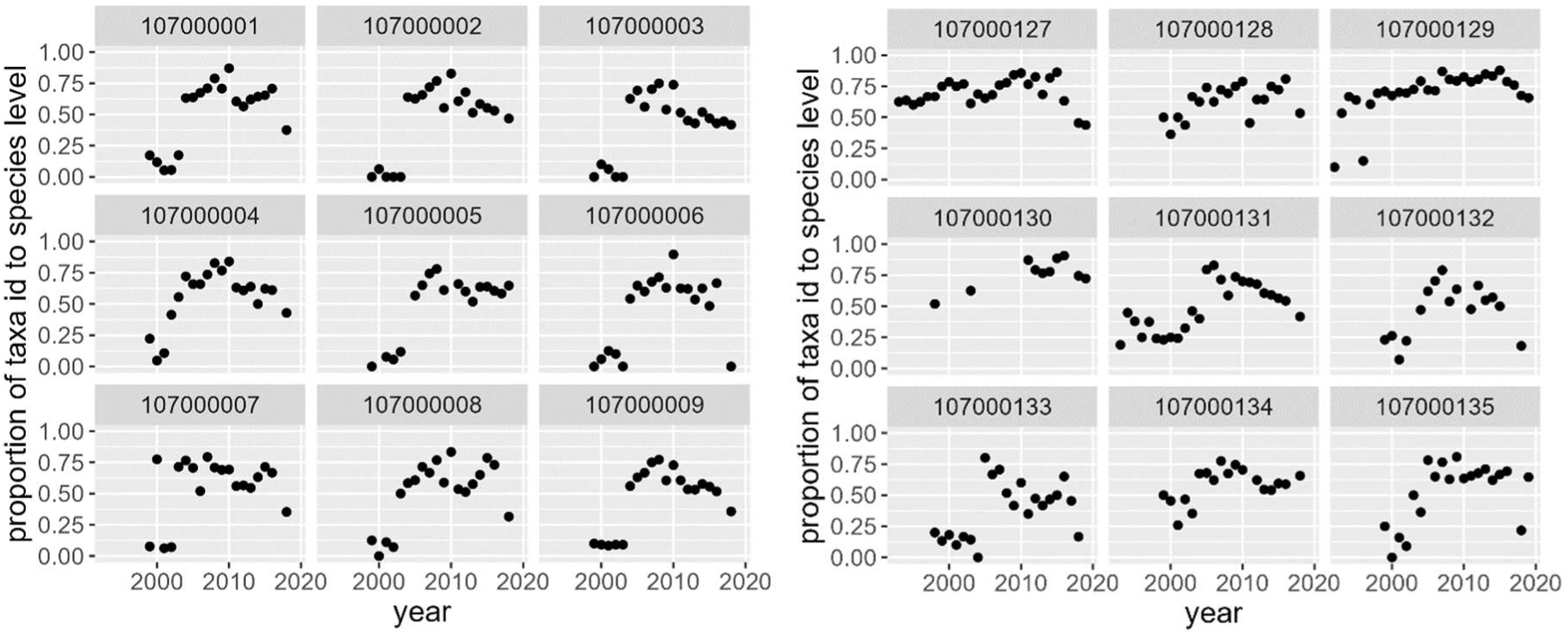
Lack of consistency in taxonomic identification in Denmark ‘species’ time series (example sites 107000001-009, 107000127-135). The y axis represents the proportion of taxa identified to species level (Fsp) within a sample.

The spatial autocorrelation of the 1816 site trend estimates was relatively high for the linear mixed effect models without study as random factor, reaching up to 0.37 (95% CI: 0.26-0.50) and 0.41 (0.28-0.61) at 10 km range for all taxa and family level, respectively (Fig. S2). When both site and study were introduced as random factor, the spatial autocorrelation of the 1816 site trend estimates was somewhat lower, reaching up to 0.20 (95% CI: 0.12-0.33) and 0.18 (0.10-0.34) for all taxa and family level (Fig. S3). Interestingly, thinning the data to a minimum distance of 20 km between sites produced the same trend estimates and p-value with only 687 sites (see Table S1).

The trend estimates for abundance at 1816 sites (period 1968-2020) were 1.01±0.01 individual per year for an estimated average abundance of 846 and 951 individuals at the centred year, with site and with/without study as random factors, respectively. The trend represented an increase of 0.12% per year and was not highly significant (*P*=0.036) when both site and study were used as random factors, and not significant (*P*=0.33) for the subset 1120 sites (period 1990-2020).

### 3.3. Taxonomic richness anomaly

Taxonomic richness anomaly was constant throughout the 1990s, rising quickly from -4 to zero at the beginning of the new millennium, before staying constant again. This pattern was similar for ‘all taxonomic ranks’ (samples with mixed taxa identified to species, genus or family) and family level, despite large changes in taxonomic resolution (proportion of taxa identified to species, genus or family) during 1990-2020 (Fig. 6). Analysis of model residuals ‘for all taxonomic ranks’ for selected year 2000 and 2010 showed significant positive spatial autocorrelations up to 0.2 in the first 50 km range (Fig. S4). This spatial autocorrelation was barely present when taxonomic data were all at family level (Fig. S5). The same analysis was repeated with the ‘species’ dataset, retaining 417 sites (Fig. 7). In this subset, both ‘all taxonomic ranks’ and family richness showed no trends from 1990 to 2020. I obtained the same results after excluding Denmark (Fig. S6) on the remaining 177 time series (with or without taxonomic corrections). The taxonomic corrections had a large effect on the Danish time series including 240 sites, effectively removing an artefact of the data (Fig. S7).

**Fig. 6.**
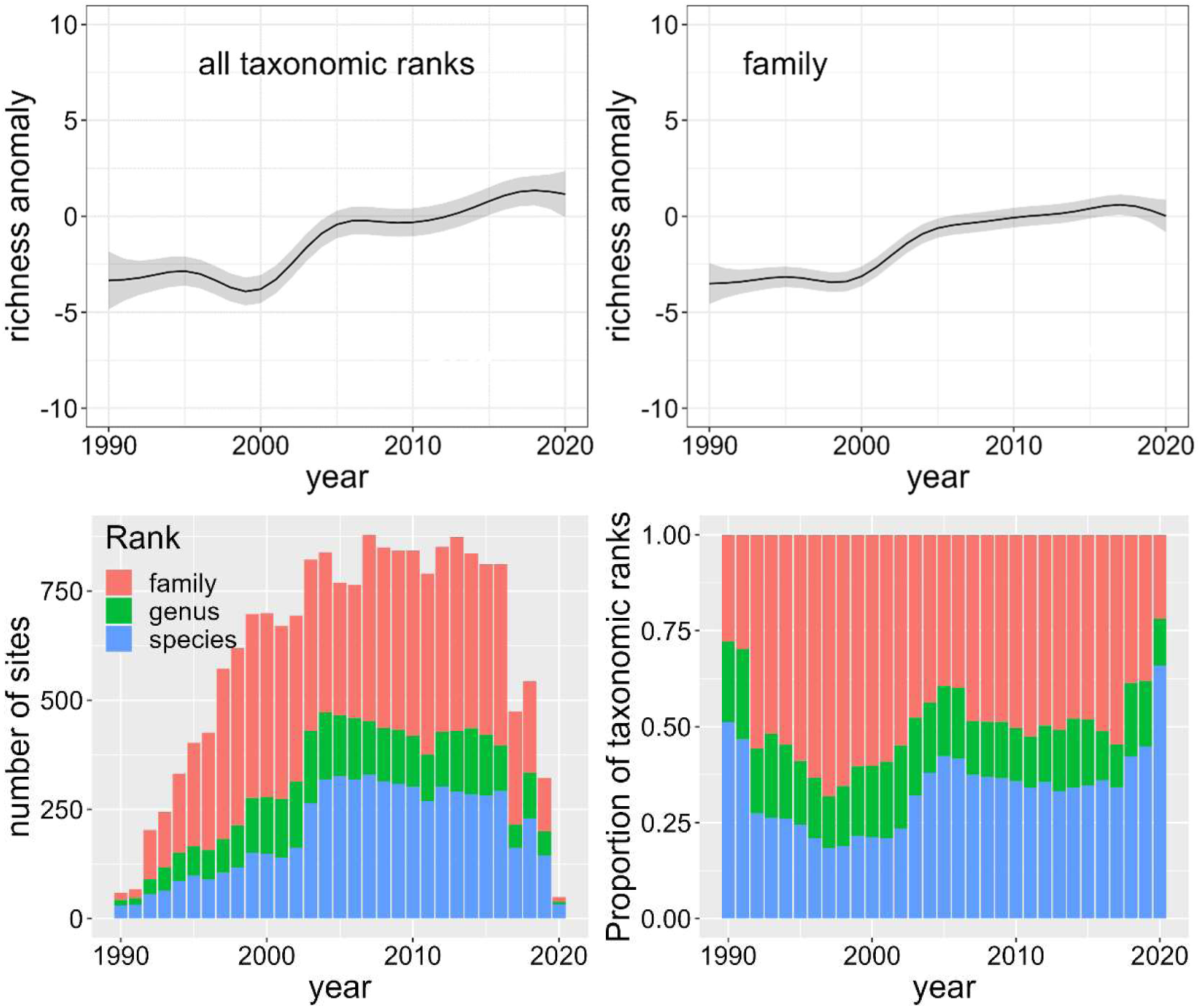
Three decades of change using all taxa at 1120 sites. Top panels: changes in taxonomic richness anomaly from long time series (gam models with 95% confidence interval) with all taxonomic ranks and family taxonomic resolution. Bottom panels: number of sites per year and proportion of taxonomic ranks in the ‘all taxonomic ranks’ dataset.

**Fig. 7.**
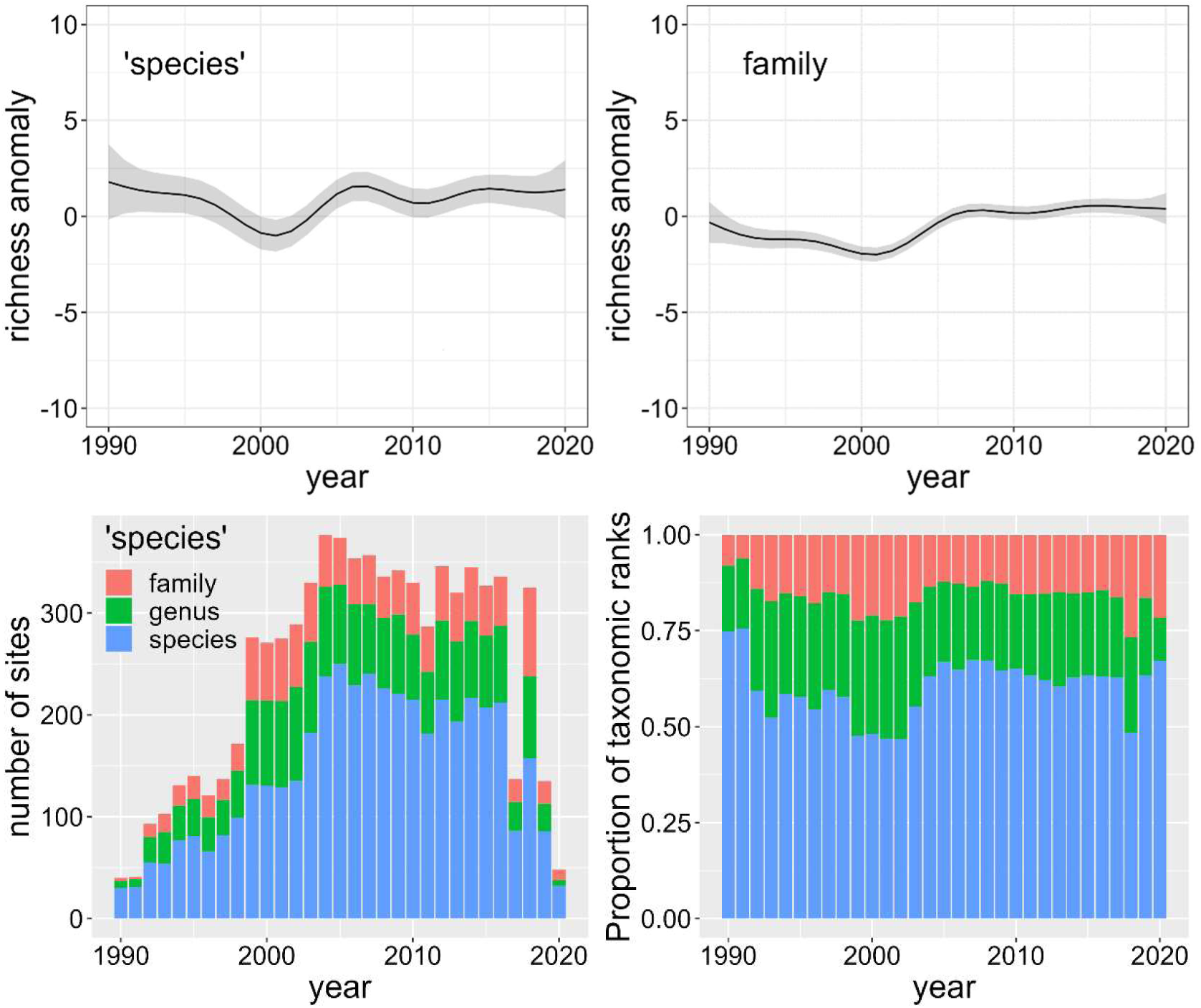
Three decades of change using ‘species’ data at 417 sites. Top panels: Changes in taxonomic richness anomaly from long time series (gam models with 95% confidence interval) with ‘species’ and family taxonomic resolution. Bottom panels: number of sites per year and proportion of taxonomic rank in the ‘species’ dataset. Note some years had less than 50% of the taxa identified to species.

Following from the above results, I plotted the density (in time) of the start and end years of the selected 1120 time series from 27 studies in the period 1990-2020 (Fig. S8). The start of each of the time series was spread throughout the 1990s and until 2004, except for a trough at the turn of the millennium. The vast majority of the times series ended post 2015. I had a closer look at the time series starting between 2001 and 2004 (inclusive), a total of 201 sites mostly (83%) from five studies (Spain_3_1, Sweden_2_1, UK_1_2, Ireland_1, Norway_3_2). The location of these times series seemed to span a wide range of environmental conditions judging from satellite imagery (Google maps). I also plotted the uncorrected median taxonomic richness (Fig. S8) and individual time series (Fig. S9) for those 27 studies. I could not see any obvious patterns of unequal representation of sites during the study period that could explain the anomaly in taxonomic richness.

The fitted changes in macroinvertebrate abundance with time was related to the observed changes in taxonomic richness for all taxonomic ranks (R^2^=0.72, *P*=0.00003) or family (R^2^=0.67, *P*=0.0001) with 1120 sites during the period 1990-2020 (*cf* Fig. 6 and Fig. 8). The fitted changes in macroinvertebrate abundance with time was also correlated to the observed small oscillations in taxonomic richness for ‘species’ (R^2^=0.49, *P*=0.0025, with some remaining temporal autocorrelation) or family (R^2^=0.53, *P*=0.0013, with some remaining temporal autocorrelation) with 417 sites during the period 1990-2020 (*cf* Fig. 7 and Fig. 8). Thus changes in taxonomic richness were related to abundance (sampling effort across sites). While the gam models explained 46% and 17 % of the deviance in abundance, respectively, the gam models explained less than 2% of the deviance in abundance anomaly, and the temporal trends were stable (oscillating) with time (Fig. 8).

**Fig. 8.**
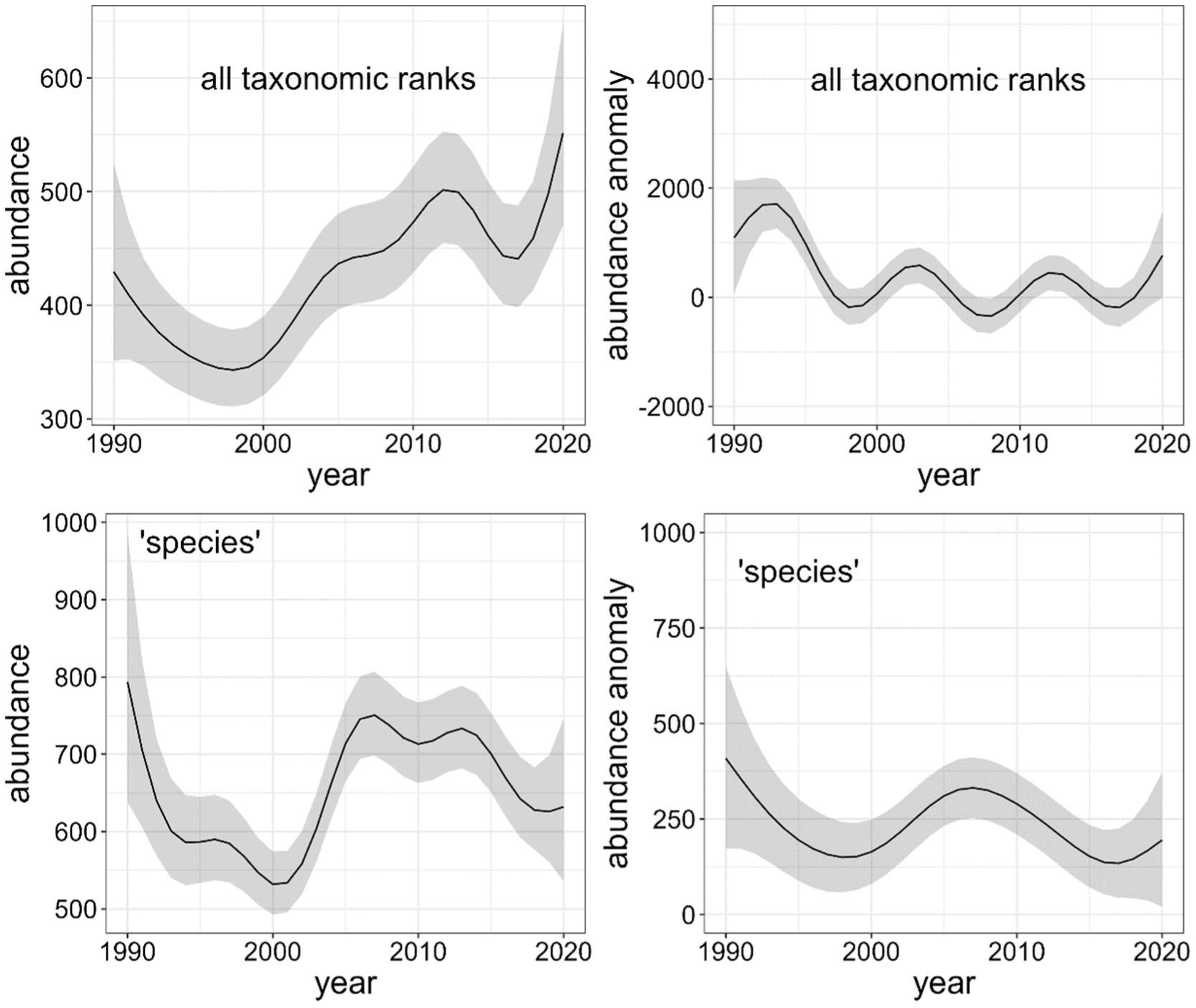
Changes in abundance (left) and abundance anomaly (right) in number of individuals, for all taxa at 1120 sites and ‘species’ data at 417 sites, for the period 1990-2020. The gam models were run on ln abundance and the fit and 95% confidence intervals were back transformed to the original scale. The partial plots were derived with latitude and longitude set at their means.

## 4. Discussion

My approach using taxonomic richness anomaly combined with corrections for intra-site taxonomic inconsistencies and overlapping long time series (at least 15 years) for the period 1990-2020 did not confirm key findings from Haase *et al*. (2023). There was no evidence of recovery in taxonomic richness during the 1990s, but instead a post-millennial short rise that may be attributed to an unequal representation of sites: a concurrent change in sampling effort across sites (abundance).

### 4.1. Trend estimates analyses using centred year

The trend estimates using linear mixed effect models returned slightly higher increase in taxa per year without (0.26±0.09) and with (0.29±0.09) taxonomic correction than Haase *et al*. (2023): 0.20 taxa per year, but were statistically not different. The coefficients from the random effects revealed high intra-site taxonomic discrepancies, in spite of Haase *et al*. (2023) fifth criterion: “We included only time series that used the same sampling method and taxonomic resolution throughout the sampling period”. The consistency of the taxonomic resolution and accuracy is often an issue when different research groups or consultants identify macroinvertebrates over many years (e.g. Haase *et al*., 2010). Such time series may not be entirely discarded, as proposed by the fifth criterion of Haase *et al*. (2023), but either reduced to family level or corrected through the model. The reduction of taxonomic resolution to family level will be necessary to assess other aspects of taxonomic diversity such as taxonomic turnover or community changes. The overall taxonomic richness increase of individual time series may also not reflect a recovery, but a systematic bias towards an earlier detection of colonisations than extinctions (Kuczynski, Ontiveros & Hillebrand, 2023). This is reinforced by the much steeper trend estimates for non-native than native species (Haase *et al*., 2023).

I am still puzzled as to why no sensitivity analyses were carried out by Haase *et al*. (2023) simply by converting all taxonomic names to family level to test the effect of taxonomic resolution. The use of their grouping ‘species’, genus (mixed) and family as fixed factor in linear mixed effect models could not be as effective as my conventional approach because the ‘species’ and genus datasets had a large proportion of taxa at a coarse taxonomic resolution (family, class). This grouping may allow the use of Integrated Distribution Models (IDMs) designed to handle datasets with varying taxonomic resolution (e.g. Adjei *et al*., 2024). I am not sure how the data in Haase *et al*., 2023 were cleaned, but a more rigorous and transparent process, such as presented in this study, must be in place to avoid intra-site discrepancies in taxonomic resolution and bias in trend estimates. Some issues of taxonomic duplication remain to be addressed, notably for species turnover and community analyses.

### 4.2. Taxonomic richness anomaly

The use of anomaly is an established approach in climate change studies, notably temperature from various proxy variables (e.g. dendrochronology). I am still reluctant to apply this approach to datasets with varying taxonomic resolutions, and I regard the results to family level as more robust (also with low spatial autocorrelation), but new statistical analyses may be able to cope with such data (Adjei *et al*., 2024). One advantage of the anomaly in taxonomic richness is to produce graphs more palatable to communicate our results and comparable to previous studies (e.g. Vaughan & Ormerod, 2012; Qu *et al*., 2023).

It is apparent that some datasets used in Haase *et al*. (2023) changed their taxonomic resolution over time within the same time series, which was then corrected in my analyses. Taxonomic richness was relatively stable over the whole period, apart from a possible sudden small rise at the turn of the millennium which was not present in the ‘species’ data subset, independently of the taxonomic resolution (all taxonomic ranks or family). Further analyses of the patterns of the time series start and end dates in the period 1990-2020 could not explain the small blip in taxonomic richness at the turn of the millennium. However, a bias arising from unequal representation of sites may be triggered by a change in sampling effort over time, not intra-site, but across sites as suggested by the concurrent changes in abundance during 1990-2020.

Haase *et al*. (2023) reported a non-significant trend estimate of 0.04 species per year for individual-based rarefied richness. Here, I refrained from reporting intra-site rarefied richness, despite the availability of easy-to-use R packages, notably iNEXT (Chao *et al*., 2014; Hsieh, Ma & Chao, 2016), due to the aggregated distribution of macroinvertebrate individuals in rivers (see assumptions in e.g. Gotelli & Colwell, 2011), the intra-site varying taxonomic resolution, and mostly the lack of changes in abundance over time within time-series, in contrast to previous studies (e.g. van Klink *et al*., 2020). Haase *et al*. (2023) claimed that their results supported previous findings by reporting a 1.17% annual increase in abundance. However, Haase *et al*. (2023) made a calculation mistake for this percentage estimate (see equationsToPercChangePerYr.xlsx in GitHub): it should be 0.14% (assuming an average abundance of 846 individuals) similar to my result of 0.12%, that is about 1 individual per year. This is a very weak trend, which is not highly significant in the present study or in Haase *et al*. (2023) for the 1816 sites in 47 studies and not significant (*P*=0.33) for the 1120 sites in 27 studies (period 1990-2020). The lack of trend was also obvious in the gam model results for abundance anomaly (period 1990-2020).

### 4.3. Further thoughts

Because of the modest but significant (short range) spatial and temporal autocorrelation in the trend estimate results and in the anomaly in taxonomic richness results, the statistical inferences (p-values) in Haase *et al*. (2023) may not be entirely reliable (Johnson *et al*., 2024). In a subsequent study, Sinclair *et al*. (2024) found no significant spatial autocorrelation in model residuals but their spatial resolution (in excess of 100 km) was too coarse. This said, I found remarkably similar results in trend estimates after pruning the dataset from 1816 to 687 sites with a minimum distance of 20 km between sites which lessen spatial autocorrelation. Both studies and countries are not independent entities and may require additional controls (Claessens, Kyritsis & Atkinson, 2023) or a more spatially explicit approach as in this study. My own study of aquatic plant time series (Demars *et al*., 2014) was also criticised for using a sub-optimal test (Kalyuzhny, 2020). These criticisms must be welcome and taken on board in future studies. A forthcoming method based on neighbourhood cross validation looks promising to deal with spatio-temporal autocorrelation in gam (Wood, 2024).

The temporal change in the anomaly of taxonomic richness during 1990-2020 were largely independent of taxonomic resolution and this may be due to the well known congruence in the taxon richness patterns of macroinvertebrates among species-, genus-, and family-level data sets (e.g. Heino & Soininen, 2007). ‘Species’ richness was indeed highly correlated to genus (mixed taxa) and family richness in Haase *et al*. (2023) data. I wonder to what extent this apparent congruence would be altered by sampling efforts. The linear mixed effect models reported that the ‘species data’ had an average taxonomic richness (intercept) of around 30 taxa per sample, barely higher than family richness (20 taxa per sample). This is tiny compared to river macroinvertebrate species richness with higher sampling efforts (see textbook Allan & Castillo, 2007, p. 233). In England and Wales, the river Wissey, Welland and Teifi (274-510 km^2^ catchment area) had far higher family (∼50), genus (∼70) and species (∼140) richness than in Haase *et al*. (2023) during a one-of survey with an intensive sample collection across river biotopes (Demars *et al*., 2012). Independently of the effect of anthropogenic impacts, current methods for biomonitoring such as 3 min kick sampling are known to harvest a small fraction of the total species richness (e.g. 50% over 18 min kick sampling in Furse *et al*., 1981) and the common practice of sub-sampling adds considerable variability in abundance and richness estimates (Clarke *et al*., 2006; Friberg *et al*., 2006). One way to lessen the lack of sampling effort per year, is to run a cumulated taxonomic richness at several temporal resolutions, i.e. summing taxa encountered over a moving window of three to five years along the time series (e.g. Demars *et al*., 2014).

More adequate and diverse methods (e.g. Malaise traps, hand net sweeping, sediment grabs) including new technology (genomics, artificial intelligence) should be used than simple kick-net or Surber sampling. The methods should be more standardised on taxonomic accumulation curves using guidance adapted to different environments. Survey must include a wider range of habitats than the traditional riffle habitat and include habitat representativity (loss, gain). The monitoring network, sampling design and protocols must be re-appraised to integrate tools such as satellite or drone imagery to meta-barcoding and meta-genomics. This will come at a cost, but can we afford not to monitor adequately biodiversity to implement policy targets (e.g. the EU’s Biodiversity Strategy for 2030)? The EU project EuropaBon has pushed hard to define Essential Biodiversity Variables (Pereira *et al*., 2013) and proposed the creation of an EU Biodiversity Observation Coordination Centre (EBOCC, Liquete *et al*., 2024). Time is of the essence (e.g. Albert *et al*., 2021).

## 5. Conclusion

Using the anomaly in taxonomic richness method showed no real change in taxonomic richness and abundance over the period 1990-2020 with samples collected across Europe. Thus, the recovery of European macroinvertebrate taxonomic richness and abundance provided by Haase *et al*. (2023) is likely to be unreliable due to biases in the characteristics of sites sampled across time. Time series analyses must be more transparent and explicit to how the data were cleaned and checked. The use of heterogeneous taxonomic data and spatio-temporal autocorrelation call for the use of alternative statistical analyses to avoid biased inferences. Finally, I question whether the current sampling efforts in European biomonitoring programmes are appropriate to monitor for biodiversity, rather than for simple indicators. Macroinvertebrates would be right to demand more from us.

## Acknowledgement

This manuscript was stimulated by discussions with David Lumsdon (soil scientist) on climate change time series analyses, and macroinvertebrate experts – some co-authors in Haase *et al*. (2023). Thank you to Joanna Lynn Kemp for reading through the manuscript and suggesting improvements prior to submission.

## Data and Code Availability Statement

Data used in this study and R scripts are available from GitHub at https://github.com/benoit-demars73/Haase_2023. Please also always refer to the original study (Haase *et al*., 2023) and published data (Welti *et al*., 2024) if using the data.

