## Supplementary information for "European freshwater macroinvertebrate richness and abundance: alternative analyses and new findings"

Benoît O.L. Demars*

Norwegian Institute for Water Research (NIVA), Økernveien 94, 0579 Oslo, Norway

**Table S1.** Effects of linear mixed-effects models on average taxonomic richness (intercept) and trend estimate (slope) on thinned dataset (20 km minimum distance between sites) reducing the number of sites from 1816 to 687. The response term Species_richness refers to the highest taxonomic resolution available in the dataset and include species, genus and families (even in the ‘species’ data). All taxonomic names were converted to family level and the analyses repeated. Note the similarity of results with Table 1.

| **Lme models** | **Intercept (se)** | **Slope (se)** | ***P*** |
|---|---|---|---|
| **Including all 687 sites in 45 studies with different taxonomic resolutions** | | | |
| Species_richness ~ cYear + (1+cYear\|site_id) | 28.7 (0.4) | 0.28 (0.02) | $<2.2e^{-16}$ |
| Species_richness ~ cYear + (1+cYear\|site_id) + (1+cYear\|study_id) | 29.3 (1.7) | 0.22 (0.08) | 0.009 |
| Species_richness ~ cYear + (1+cYear\|site_id) + (1+cYear\|c.Fy_id_round) | 24.3 (1.6) | 0.31 (0.03) | 0.0003 |
| Species_richness ~ cYear + (1+cYear\|site_id) + (1+cYear\|study_id) + (1+cYear\|c.Fy_id_round) | 24.6 (2.3) | 0.29 (0.08) | 0.003 |
| **Including all 687 sites in 45 studies at family taxonomic resolution** | | | |
| Family_richness ~ cYear + (1+cYear\|site_id) | 23.2 (0.3) | 0.20 (0.02) | $<2.2e^{-16}$ |
| Family_richness ~ cYear + (1+cYear\|site_id) + (1+cYear\|study_id) | 21.3 (1.0) | 0.15 (0.03) | 0.0007 |

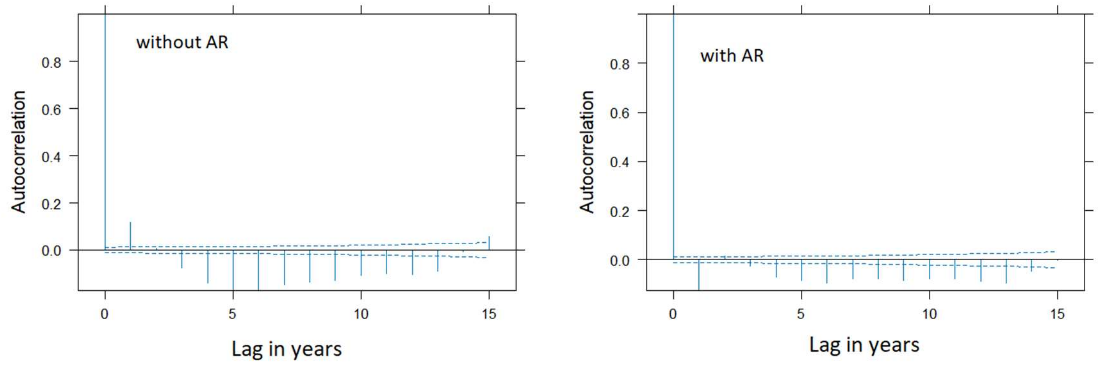


Fig. S1. Temporal autocorrelation with/without an autoregressive term (AR) in linear mixed effect model testing the effect of cYear on taxonomic richness at 1816 sites with site as random effect (slope and intercept). The same results were obtained when taxonomic richness was corrected with c.Fy (see middle panel of Fig. 4) prior to running the model. R script available in Supplementary information.

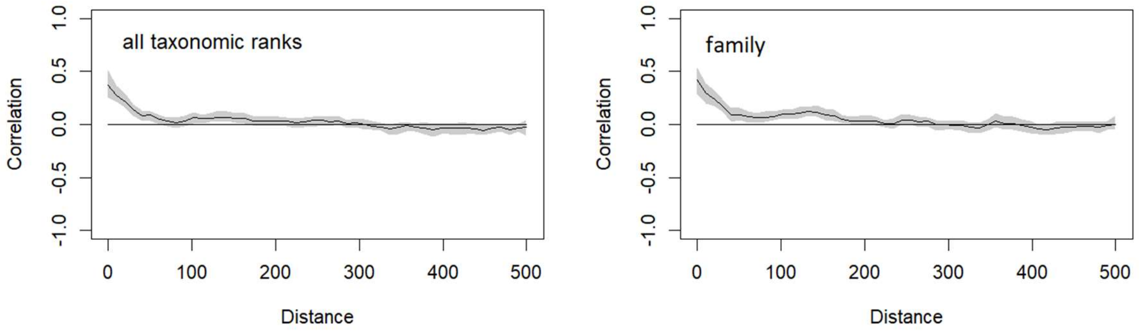


Fig. S2. Spatial autocorrelation in 1816 site trend estimates from linear mixed effect models *with site* as random factor for all taxonomic ranks (with an additional random factor for taxonomic correction) and family level. Distance is in km.

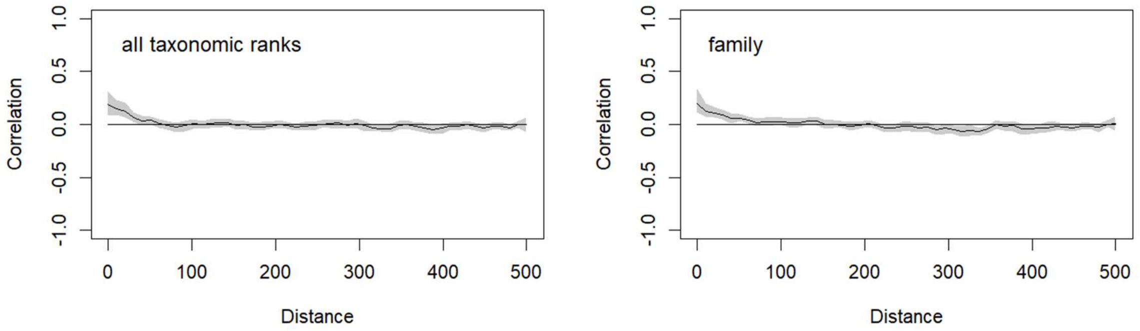


Fig. S3. Spatial autocorrelation in 1816 site trend estimates from linear mixed effect models *with site and study* as random factors for all taxonomic ranks (with an additional random factor for taxonomic correction) and family level. Distance is in km.

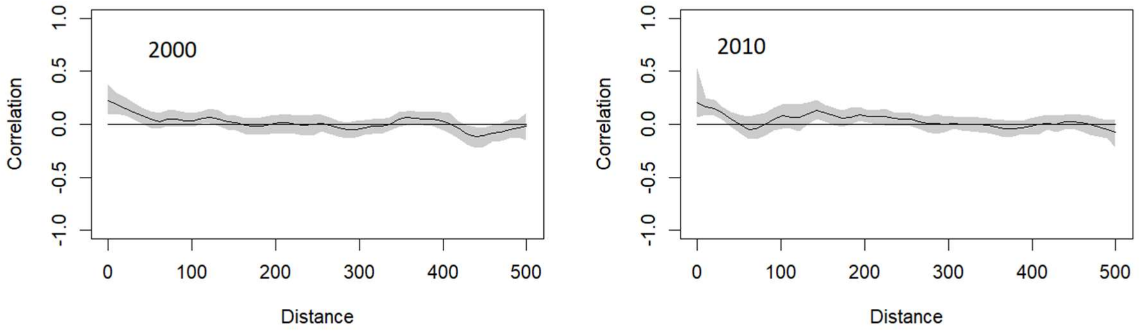


Fig. S4. Mixed taxa (species, genus and family): Spatial autocorrelation in gam model response residuals (taxonomic richness) from selected year 2000 (699 sites) and year 2010 (843 sites). Distance is in km.

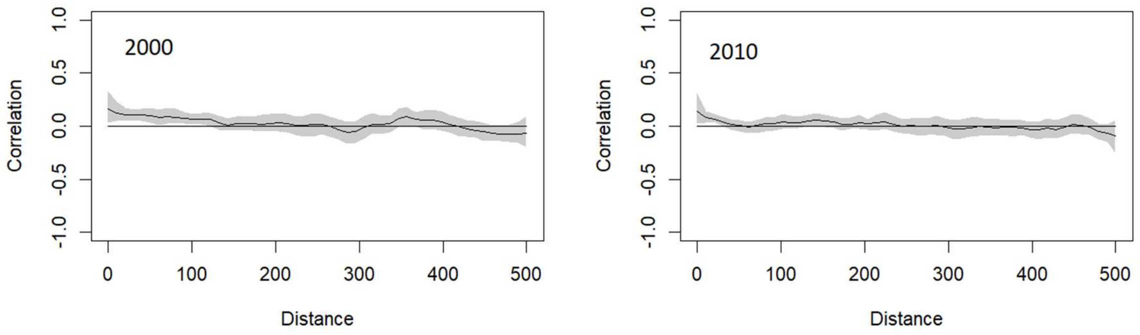


Fig. S5. Family: Spatial autocorrelation in gam model response residuals (taxonomic richness) from selected year 2000 (699 sites) and year 2010 (843 sites). Distance is in km.

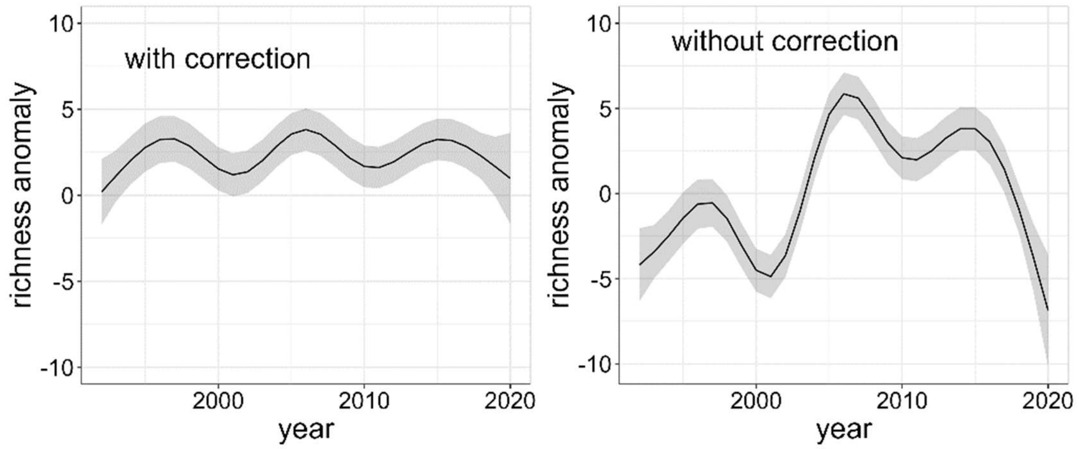


Fig. S6. Effect of taxonomic correction on richness anomaly trend in Denmark. Changes (gam models with 95% confidence interval) in taxonomic richness anomaly over three decades using 'species' data and 240 long time series (range ≥15 years, records ≥ 8 sampling years) with and without a correction for taxonomic resolution.

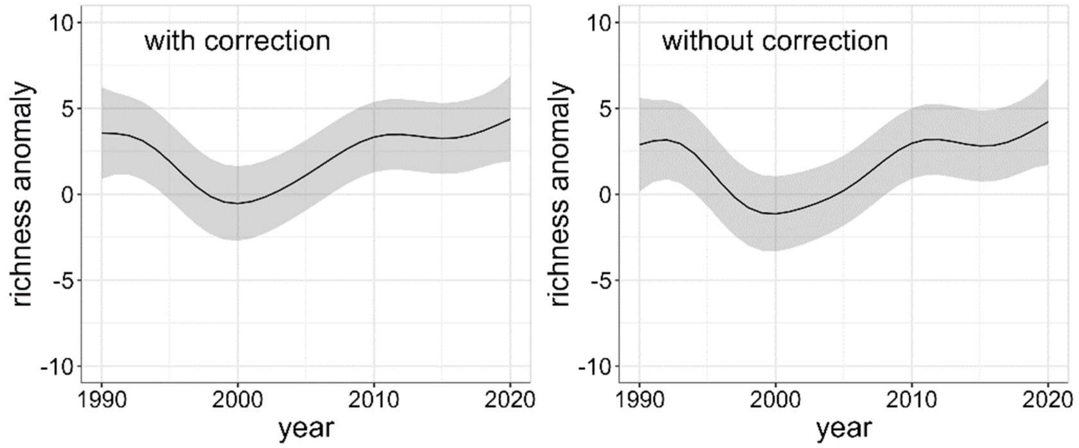


Fig. S7. Taxonomic correction had no effect on richness anomaly trend for time series with consistent taxonomic identification. Changes (gam models with 95% confidence interval) in taxonomic richness anomaly over three decades using 'species' data and 177 long time series (range ≥15 years, records ≥ 8 sampling years) with and without a correction for taxonomic resolution.

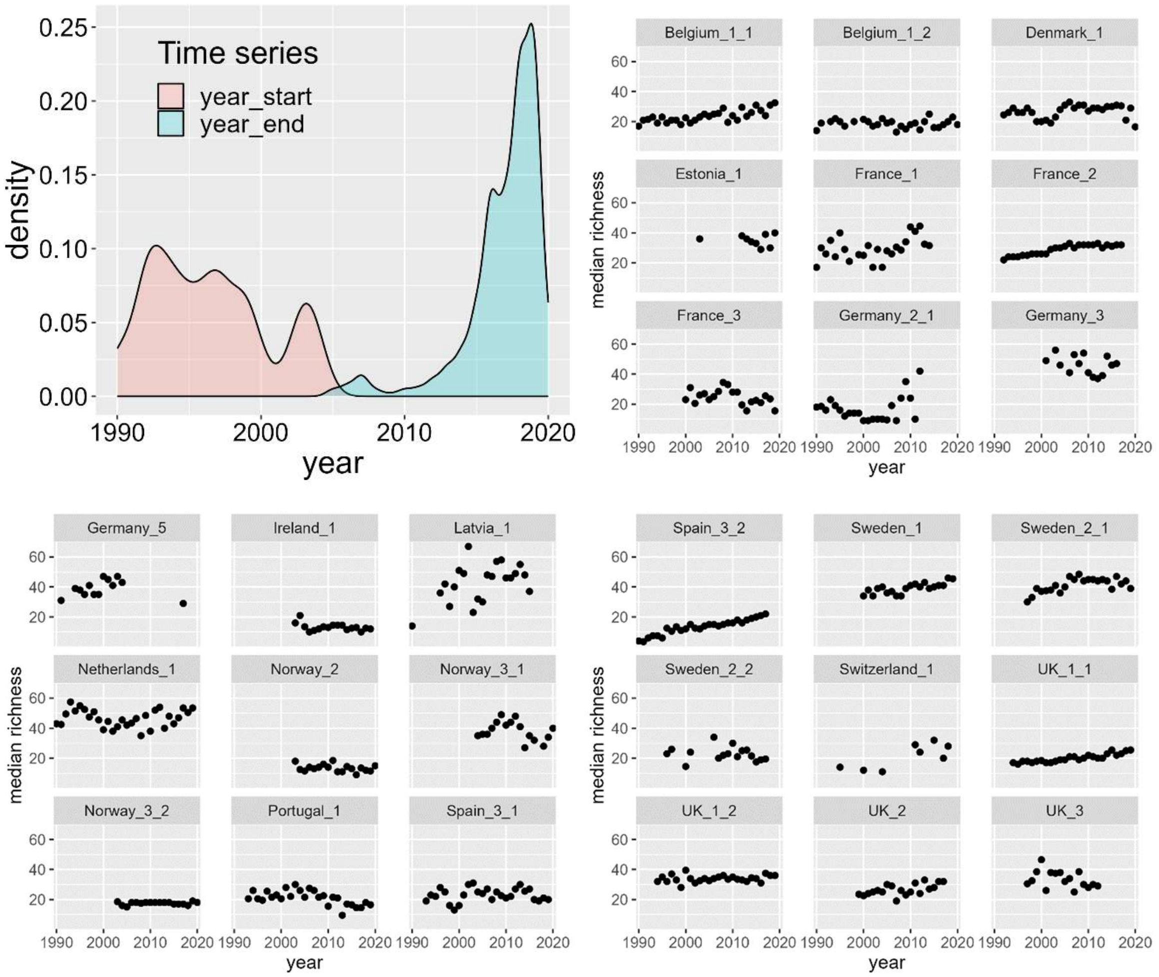


Fig. S8. Density in the start and end year for 1120 time series in 27 studies, and uncorrected median taxonomic richness for the 27 studies, for the period 1990-2020. Note the number of sites can vary a lot within study over time, so these graphs were just used for a first look.

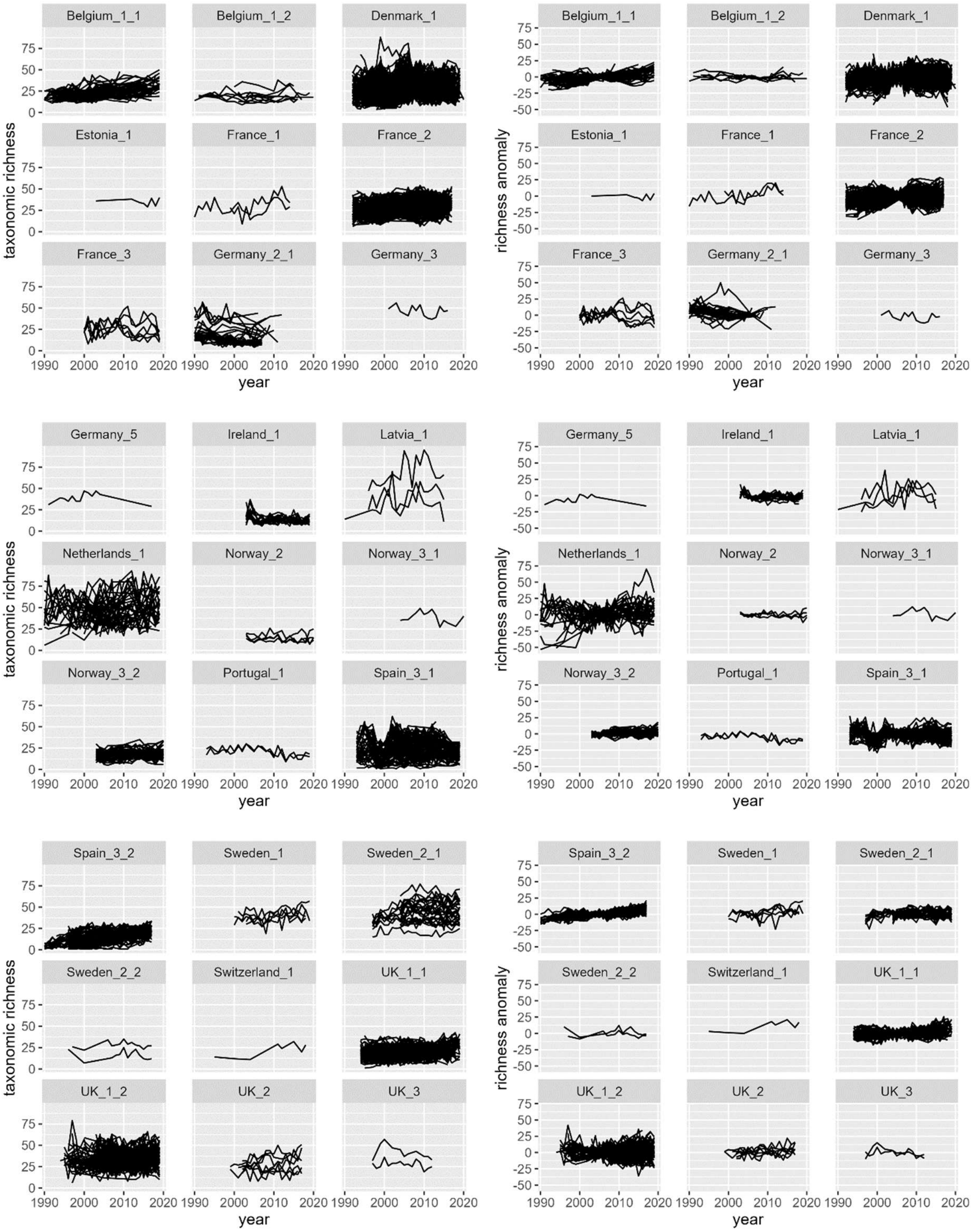


Fig. S9. Taxonomic richness (left) and anomaly in taxonomic richness (right) for 1120 time series in 27 studies for the period 1990-2020.

Selection of GAM model results

All taxonomic ranks 1990-2020: summary results of gam model with 1120 sites (Fig. 6)

```
Family: gaussian
Link function: identity

Formula:
Species_richness_5y ~ s(year) + s(Lon, Lat) + s(c.Fy_id)

Parametric coefficients:
            Estimate Std. Error t value Pr(>|t|)
(Intercept) -0.74511    0.05597  -13.31   <2e-16 ***
---
Signif. codes:  0 '***' 0.001 '**' 0.01 '*' 0.05 '.' 0.1 ' ' 1

Approximate significance of smooth terms:
              edf Ref.df      F p-value
s(year)     7.924  8.682  92.08  <2e-16 ***
s(Lon,Lat) 28.309 28.966  39.33  <2e-16 ***
s(c.Fy_id)  8.149  8.814 157.70  <2e-16 ***
---
Signif. codes:  0 '***' 0.001 '**' 0.01 '*' 0.05 '.' 0.1 ' ' 1

R-sq.(adj) =  0.169   Deviance explained = 17.1%
-REML =  64515  Scale est. = 58.442    n = 18657
```

Family 1990-2020: summary results of gam model with 1120 sites (Fig. 6)

```
Family: gaussian
Link function: identity

Formula:
Family_richness_5y ~ s(year) + s(Lon, Lat)

Parametric coefficients:
            Estimate Std. Error t value Pr(>|t|)
(Intercept) -0.32877    0.04043  -8.132 4.49e-16 ***
---
Signif. codes:  0 '***' 0.001 '**' 0.01 '*' 0.05 '.' 0.1 ' ' 1

Approximate significance of smooth terms:
              edf Ref.df      F p-value
s(year)     7.736  8.574 136.75  <2e-16 ***
s(Lon,Lat) 28.281 28.963  41.24  <2e-16 ***
---
Signif. codes:  0 '***' 0.001 '**' 0.01 '*' 0.05 '.' 0.1 ' ' 1

R-sq.(adj) =  0.108   Deviance explained = 10.9%
-REML =  58433  Scale est. = 30.496    n = 18657
```

'Species' 1990-2020: summary results of gam model with 417 sites (Fig. 7)

```
Family: gaussian
Link function: identity

Formula:
Species_richness_5y ~ s(year) + s(Lon, Lat) + s(c.Fsp_id)

Parametric coefficients:
            Estimate Std. Error t value Pr(>|t|)
(Intercept) -0.96964    0.09558  -10.14   <2e-16 ***
---
Signif. codes:  0 '***' 0.001 '**' 0.01 '*' 0.05 '.' 0.1 ' ' 1

Approximate significance of smooth terms:
               edf Ref.df       F p-value
s(year)      7.309  8.292   7.371  <2e-16 ***
s(Lon,Lat)  26.673 28.642  23.290  <2e-16 ***
s(c.Fsp_id)  7.195  8.258 139.980  <2e-16 ***
---
Signif. codes:  0 '***' 0.001 '**' 0.01 '*' 0.05 '.' 0.1 ' ' 1

R-sq.(adj) =  0.255   Deviance explained = 25.9%
-REML =  26541  Scale est. = 68.473    n = 7495
```

Family 1990-2020: summary results of gam model with 417 sites (Fig. 7)

```
Family: gaussian
Link function: identity

Formula:
Family_richness_5y ~ s(year) + s(Lon, Lat)

Parametric coefficients:
            Estimate Std. Error t value Pr(>|t|)
(Intercept)  0.05757    0.05193   1.109    0.268

Approximate significance of smooth terms:
              edf Ref.df     F p-value
s(year)     7.447  8.388 32.93  <2e-16 ***
s(Lon,Lat) 26.072 28.444 19.96  <2e-16 ***
---
Signif. codes:  0 '***' 0.001 '**' 0.01 '*' 0.05 '.' 0.1 ' ' 1

R-sq.(adj) =  0.101   Deviance explained = 10.5%
-REML =  21958  Scale est. = 20.212    n = 7495
```

Abundance 1990-2020: summary results of gam model with 1120 sites (Fig. 8)

```
Family: gaussian
Link function: identity

Formula:
log(abundance) ~ s(year) + s(Lon, Lat)

Parametric coefficients:
            Estimate Std. Error t value Pr(>|t|)
(Intercept) 6.940927   0.007684   903.3   <2e-16 ***
---
Signif. codes:  0 '***' 0.001 '**' 0.01 '*' 0.05 '.' 0.1 ' ' 1

Approximate significance of smooth terms:
              edf Ref.df      F p-value
s(year)     7.635  8.511  29.51  <2e-16 ***
s(Lon,Lat) 28.735 28.995 540.93  <2e-16 ***
---
Signif. codes:  0 '***' 0.001 '**' 0.01 '*' 0.05 '.' 0.1 ' ' 1

R-sq.(adj) =  0.459   Deviance explained = 46.1%
-REML =  27469  Scale est. = 1.1014    n = 18655
```

Abundance anomaly 1990-2020: summary results of gam model with 1120 sites (Fig. 8)

```
Family: gaussian
Link function: identity

Formula:
log(abundance_anomaly) ~ s(year) + s(Lon, Lat)

Parametric coefficients:
             Estimate Std. Error t value Pr(>|t|)
(Intercept) 10.530310   0.001078    9767   <2e-16 ***
---
Signif. codes:  0 '***' 0.001 '**' 0.01 '*' 0.05 '.' 0.1 ' ' 1

Approximate significance of smooth terms:
              edf Ref.df      F p-value
s(year)     8.268  8.844 13.631  <2e-16 ***
s(Lon,Lat) 11.399 15.327  8.264  <2e-16 ***
---
Signif. codes:  0 '***' 0.001 '**' 0.01 '*' 0.05 '.' 0.1 ' ' 1

R-sq.(adj) =  0.0131   Deviance explained = 1.41%
-REML = -9219.7  Scale est. = 0.021686  n = 18655
```

Abundance 1990-2020: summary results of gam model with 417 sites (Fig. 8)

```
Family: gaussian
Link function: identity

Formula:
log(abundance) ~ s(year) + s(Lon, Lat)

Parametric coefficients:
            Estimate Std. Error t value Pr(>|t|)
(Intercept)  6.47273    0.01067   606.5   <2e-16 ***
---
Signif. codes:  0 '***' 0.001 '**' 0.01 '*' 0.05 '.' 0.1 ' ' 1

Approximate significance of smooth terms:
              edf Ref.df     F p-value
s(year)     7.514  8.434 13.75  <2e-16 ***
s(Lon,Lat) 26.590 28.617 49.52  <2e-16 ***
---
Signif. codes:  0 '***' 0.001 '**' 0.01 '*' 0.05 '.' 0.1 ' ' 1

R-sq.(adj) =  0.168   Deviance explained = 17.2%
-REML =  10101  Scale est. = 0.85323   n = 7491
```

Abundance anomaly 1990-2020: summary results of gam model with 417 sites (Fig. 8)

```
Family: gaussian
Link function: identity

Formula:
log(abundance_anomaly) ~ s(year) + s(Lon, Lat)

Parametric coefficients:
            Estimate Std. Error t value Pr(>|t|)
(Intercept) 9.458273   0.001181    8006   <2e-16 ***
---
Signif. codes:  0 '***' 0.001 '**' 0.01 '*' 0.05 '.' 0.1 ' ' 1

Approximate significance of smooth terms:
              edf Ref.df     F  p-value
s(year)     4.989  6.085 3.931 0.000581 ***
s(Lon,Lat) 15.395 19.964 3.279 8.21e-07 ***
---
Signif. codes:  0 '***' 0.001 '**' 0.01 '*' 0.05 '.' 0.1 ' ' 1

R-sq.(adj) =  0.0121   Deviance explained = 1.47%
-REML = -6412.4  Scale est. = 0.010455  n = 7491
```
